# HSV-1 hijacks the DNA repair protein RAD51 at gene promoters to drive viral transcription

**DOI:** 10.64898/2026.07.09.737117

**Authors:** Namrata Kumar, Laura E. M. Dunn, Tanner M. Tessier, Katharina E. Hayer, Sireen Sweed, Maya Ralph-Altman, Holly M. Chan, Brandon G. Waxman, Edwin Halko, Yuval Altman, Jonathan Korner, Oren Kobiler, Joel D. Baines, Matthew D. Weitzman

**Affiliations:** Division of Protective Immunity, and Division of Cancer Pathobiology, Department of Pathology and Laboratory Medicine, The Children’s Hospital of Philadelphia, Philadelphia, PA, USA; Department of Pathology and Laboratory Medicine, Perelman School of Medicine, University of Pennsylvania, Philadelphia, PA, USA; Baker Institute for Animal Health, College of Veterinary Medicine, Cornell University, Ithaca, New York, USA; Department of Biomedical and Health Informatics, Children’s Hospital of Philadelphia, Philadelphia, USA; Department of Microbiology and Infectious diseases, Gray School of Medical Sciences, Gray Faculty of Medical and Health Sciences, Tel Aviv University, Tel Aviv, Israel; Tzur Yitzhak, Israel; Modi’in-Maccabim-Re’ut, Israel; Penn Epigenetics Institute, Perelman School of Medicine, University of Pennsylvania, Philadelphia, PA, USA

## Abstract

Productive infection by Herpes Simplex Virus type 1 (HSV-1) requires initiation of efficient viral gene expression. Upon nuclear entry, HSV-1 genomes are associated with several cellular factors, including DNA repair proteins. It is unclear how these cellular factors impact viral processes at early stages of infection. In this study, we investigate the role of RAD51, a core homologous recombination (HR) factor. We measured nascent viral transcription and show that RAD51 plays a pro-viral role in infection by promoting immediate-early viral gene expression. We demonstrate that RAD51 binds GC-rich gene promoter of ICP4 and directly promotes gene expression. We also reveal a previously unknown interaction between RAD51 and ADNP, a subunit of the ChAHP complex, known for its role in transcription regulation. We propose a model where RAD51 binds incoming genomes at promoter regions regulating the genome landscape and allowing for efficient transcription initiation.

## Introduction

Maintenance of genome stability is essential for cellular function and survival. To deal with DNA damage, cells have evolved specific DNA repair pathways (1–4). Homologous recombination (HR) is one highly conserved DNA repair pathway that repair double-stranded breaks (DSBs) and interstrand crosslinks (ICLs) (5–7). DSBs are one of the most toxic DNA lesions and can arise from exogenous sources, such as radiation and chemotherapy, as well as from endogenous sources, such as reactive oxygen species, metabolic byproducts, and replication stress. If left unrepaired, DSBs are lethal and can lead to genome instability, genomic rearrangements, and cancer(5,8–14).

RAD51 is a core factor in HR, responsible for finding and invading the undamaged homologous sequence for error-free repair of DSBs (5,10,15–17). It is an ATPase that forms nucleofilaments on single-stranded DNA with high affinity. Even though RAD51 is essential for HR, unlike other HR factors, very few mutations in RAD51 are implicated in cancer predisposition. In contrast, overexpression of RAD51 is common in various cancer types (10,12,14,18–20). This phenomenon called the ‘RAD51 paradox’ distinguishes RAD51 from other HR factors and suggests that RAD51 may have roles that are distinct from HR. A well-studied HR-independent role for RAD51 is in DNA fork protection and reversal (16). More recently, RAD51 has been shown to associate with DNA secondary structures such as DNA:RNA hybrids (21–23) and G-quadruplexes (24), however the mechanism and implications of these interactions are still under investigation. While the role of RAD51 is well studied in the context of DSBs, new roles are still emerging and being explored.

Viral DNA genomes present the cell with large amounts of exogenous genetic material with unusual DNA structures, which challenge the host genome and recruit host factors for viral benefit (25–28). HSV-1 is a double-stranded DNA (dsDNA) virus of 152kbp and replicates in the host cell nucleus during infection. Upon releasing its dsDNA genome in the host cell nucleus, HSV-1 undergoes a sequential cascade of gene expression. Viral and cellular proteins regulate the transcription, replication, and packaging of viral genomes. HSV-1 codes for approximately 80 genes, which are expressed in temporally regulated phases: immediate-early (IE or α), early (E or β) and late (L or γ) (29–31). HSV-1 gene expression is mediated by the cellular RNA polymerase II (RNA Pol II) and other cellular proteins, with help from viral transcription factors, VP16 and ICP4. Upon nuclear entry, the viral tegument protein VP16 directly binds IE gene promoters to mediate immediate-early gene expression, including ICP4 (32). ICP4 stimulates expression of E and L genes, including the genes encoding for DNA replication. Expression of L genes is dependent on the onset of viral DNA replication (33–35).

It is well established that virus infection requires manipulation of cellular processes, including the host DNA replication and DNA damage response (DDR) machinery (33,36–44). Many cellular DNA replication and repair proteins are enriched on viral genomes during HSV-1 infection. In some cases, DNA repair proteins are beneficial for infection (37,39). On the other hand, there is also selective targeting of DNA repair proteins through ubiquitylation by the HSV-encoded E3 ubiquitin ligase ICP0 (45,46). Recent studies have shown that DDR activation during HSV-1 infection is distinct from canonical DDR signaling and independent of DNA damage (47,48).

RAD51 is one of many cellular DNA repair factors previously reported to be recruited to HSV-1 Replication Compartments (42), although the role of RAD51 during HSV-1 infection is not understood. Here we employ herpes simplex virus type 1 (HSV-1) infection as a model system to investigate the role of RAD51 during virus infection. We found that loss or inhibition of RAD51 caused a significant defect in producing infectious viral progeny and viral proteins. This defect was mediated by a decrease in immediate-early viral transcription when RAD51 was inhibited. We found that RAD51 binds a GC-rich viral promoter and directly drives gene expression. We demonstrate that RAD51 is independent of its known role in recombination since RAD51 inhibition has no effect on viral intergenomic recombination. Finally, we revealed a previously unknown interaction of RAD51 with ADNP, a core factor of the ChAHP complex, known to restrict chromatin accessibility and transcription repression. Together, our data support a model where RAD51 recognizes specific regions on viral gene promoters, potentially regulating the transcriptional landscape to drive efficient gene expression and productive infection.

## Results

### RAD51 loss leads to decreased HSV-1 progeny and protein production

To assess the extent to which RAD51 has a functional role in infection, we knocked down RAD51 using siRNA or inhibited its function using commercially available inhibitors in Human Foreskin Fibroblasts (HFFs). We then infected depleted cells with HSV-1 and tested how RAD51 loss impacted viral progeny production (**Figure 1A**). We found that loss of RAD51 function led to a 10-20-fold decrease in progeny production at both 24 and 48 hours post infection (hpi). As a complementary approach, we inhibited RAD51 function by using commercially available RAD51 inhibitors, B02, RI-1, RI-2, and IBR2 **(Figure 1B)**, and observed a similar decrease in progeny production. To assess whether RAD51 affected viral replication, we visualized VRCs through immunofluorescence for ICP8, the single-stranded DNA binding encoded by the virus (49,50). We categorized VRCs into early, early-mid, late-mid, and late structures and analyzed the number of cells in each stage of infection at 4, 6 and 8 hpi. We noted that in the absence of RAD51, progression of VRCs to later stages of infection was delayed or limited (**Figure 1C**), suggesting that RAD51 is essential for progression of HSV-1 infection. We next examined viral protein levels during infection to determine which stage (IE, E or L) was impacted by RAD51 loss (**Figure 1D**, **Supplementary Figure S1A**). When compared to controls, we found that RAD51-deficient cells produced significantly lower levels of viral proteins at all stages, including immediate-early (IE), early (E) and late (L) viral proteins. RAD51 can protect DNA replication forks during replication stress (16). To assess whether RAD51 was involved in preventing viral DNA replication stress, we treated infected cells with two viral DNA replication inhibitors, phosphonoacetic acid (PAA) or Pritelivir (**Figure 1E**, **Supplementary Figure S1B**). This inhibitor prevents expression of late genes and proteins (see VP21). If the defect observed in RAD51-deficient cells were due to its role in mediating viral DNA replication, we would not observe a change in protein levels in the absence of viral DNA replication. However, we still noted lower protein levels in RAD51-deficient cells **(Figure 1E).** Together, these data indicate that RAD51 plays a role in HSV-1 infection during immediate-early stages of infection prior to the onset of viral DNA replication.

**Figure 1:**
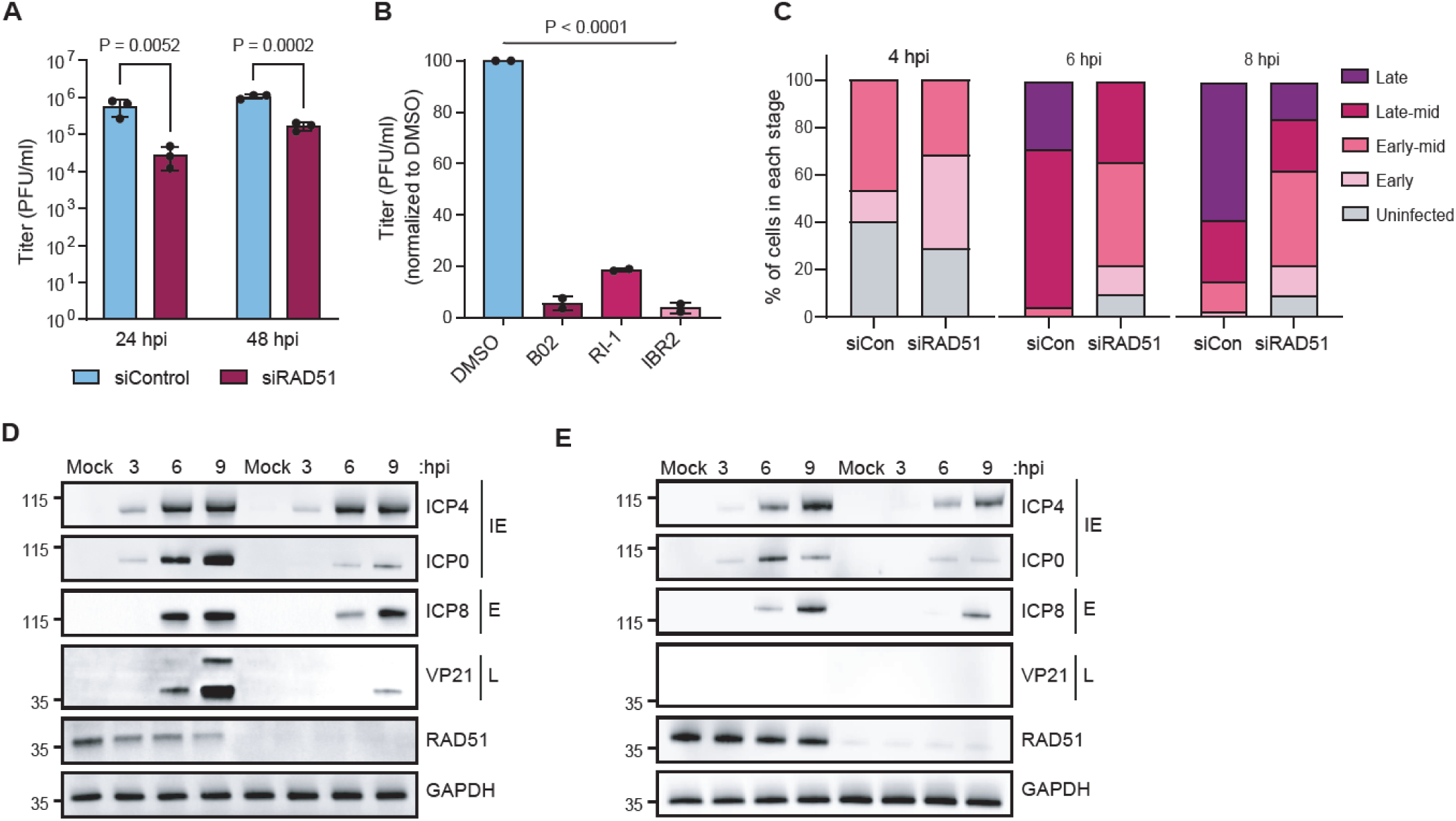
RAD51 loss leads to decreased HSV-1 progeny and protein production. **A.** HFF cells were transfected with a control or RAD51 siRNA. 48 hours post transfection, cells were infected with HSV-1 17syn+ at MOI 0.1 and harvested for plaque assay at 24 and 48 hours post infection (hpi). Plaque assay was performed in Vero cells and plaques were counted after 3-4 days. P-values derived from Two-Way ANOVA. **B.** HFF cells were pre-treated with DMSO or indicated RAD51 inhibitors for 16 hours. Cells were then infected with HSV-1 17syn+ at MOI 0.1 and harvested for plaque assay at 24 hpi. Plaque assay was performed in Vero cells and plaques were counted after 3-4 days. P-values derived from One-Way ANOVA. **C.** HFF cells were transfected with a control or RAD51 siRNA. 48 hours post transfection, cells were infected with HSV-1 17syn+ at MOI 3 and fixed at 4, 6 and 8 hpi for immunofluorescence. Antibody against ICP8 was used to mark viral replication compartments (VRCs) and quantified based on the stage of infection. **D.** HFF cells were transfected with a control or RAD51 siRNA. 48 hours post transfection, cells were infected with HSV-1 17syn+ at MOI 3 and harvested for western blot at 3, 6, 9 hpi**. E.** Cells were treated the same as D. Phosphonoacetic acid (PAA) was added at 1 hpi.

### The role of RAD51 during infection is distinct from functions in DNA replication and recombination

We found that RAD51-deficient cells were defective in early viral protein production (**Figure 1E**). We therefore analyzed the recruitment of RAD51 to VRCs during infection in the absence of viral DNA replication (**Figure 2A, 2B**). We treated cells with PAA and performed immunofluorescence and examined the colocalization of RAD51 with ICP8. We found that RAD51 colocalization with ICP8 was unaffected when viral DNA replication was inhibited. In contrast, when we performed immunofluorescence for another core HR protein, RPA, we found that the recruitment to VRCs decreased when viral DNA replication was inhibited (**Figure 2C, 2D**). Additionally, knocking down RPA using siRNA showed a milder (2-3-fold) decrease in viral progeny production and viral protein levels compared to RAD51 loss/ inhibition (**Supplementary Figure S2A, S2B**), and this was further unaffected when cells were treated with PAA (**Supplementary Figure S2C)**. On the other hand, cells lacking BRCA2, the obligate loading factor for RAD51, also displayed a 10-fold decrease in infectious progeny production, consistent with published studies (37) (**Supplementary Figure S2D**). This finding highlights a key difference in canonical HR and the functions of RAD51 in response to HSV-1 infection (**Figure 2E**).

**Figure 2:**
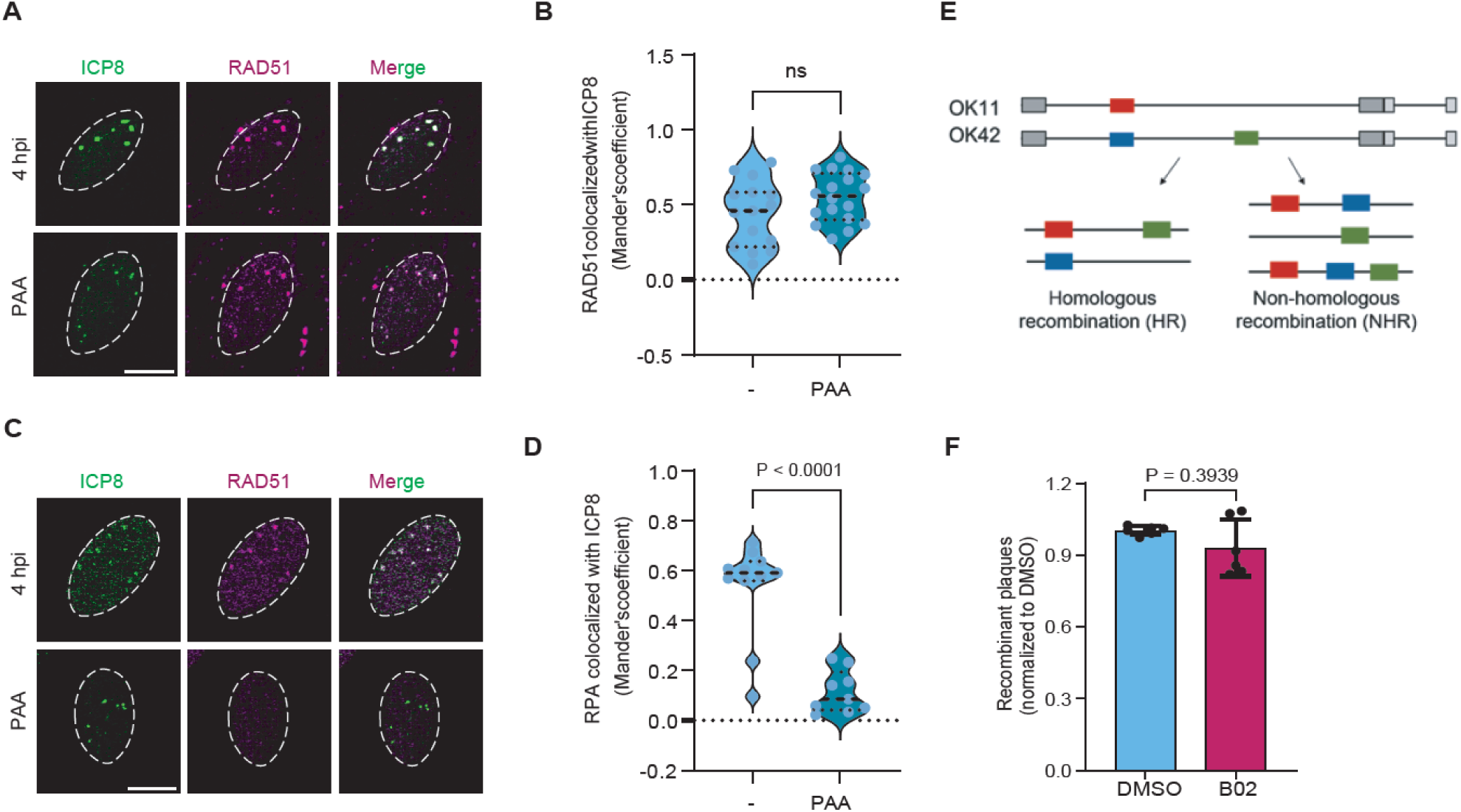
The role of RAD51 during infection is distinct from functions in DNA replication and recombination. **A.** HFF cells were infected with HSV-1 17syn+ at MOI 3 and fixed at 4 hpi for immunofluorescence. **B**. Antibodies against RAD51 and ICP8 were used to quantify colocalization at VRCs**. C.** HFF cells were infected with HSV-1 17syn+ at MOI 3 and fixed at 4 hpi for immunofluorescence**. D.** Antibodies against RPA and ICP8 were used to quantify colocalization at VRCs. **E.** Schematic showing viruses used for the viral intergenomic recombination assay. F. HFF cells were treated with DMSO or 20uM B02 for 2 hours pre-infection. Cells were co-infected with OK11 and OK42 at MOI 20 and homology-mediated recombinant plaques were quantified.

RAD51 loss reduces HR efficiency (51,52). Several groups have used elegant GFP reporter systems to study recombination and examine changes when HR factors are inhibited (53–55). These systems were employed within the cellular genome during HSV-1 infection to demonstrate that while recombination via HR and NHEJ were inhibited, the Single Strand Annealing (SSA) pathway was activated (56). The authors concluded that SSA is the homology-mediated repair pathway utilized during HSV-1 infection, while HR was inhibited. To assess whether RAD51 inhibition directly changes viral genome recombination, we utilized a reporter system directly integrated into the virus. Two different HSV-1 viruses, OK11 expressing the mCherry (RFP) gene located between UL37 and UL38 viral genes, and OK42 expressing both mTurq2 (CFP) and mVenus (YFP) genes located between UL37 and UL38 genes and fused within the UL25 gene, respectively (57) (**Figure 2e**). These viruses were co-infected at high multiplicity of infections (MOI 20) to induce intergenomic recombination, and progeny were examined by plaque assay. Based on the color of the plaques, we can determine recombination events on the viral genomes. Using this approach, we showed that RAD51 inhibition had no impact on intergenomic recombination events (**Figure 2f**), indicating that the role of RAD51 during infection is distinct from its role in HR.

### RAD51 inhibition leads to decreased nascent viral transcription

Upon nuclear entry of the viral dsDNA, HSV-1 IE genes are expressed (44). To evaluate whether RAD51 plays a role in mediating viral transcription, we treated HFF cells with or without B02 for 2 hours pre-infection. Single-molecule RNA FISH was performed to visualize ICP4 and ICP8 viral mRNAs at 3 hpi. We observed that RAD51 inhibition led to a significant 2-3-fold decrease in mRNA expression **(Figure 3A-C)**. To examine viral transcription at a single-molecule level, we treated HEp2 cells with DMSO or the B02 inhibitor and employed Precision Run-On Sequencing (PRO-seq). PRO-seq is a nascent-transcription mapping technique that reveals RNAPII locations on DNA at a single-molecule resolution (58). By measuring actively engaged RNAPII, PRO-seq analysis can identify promoter-proximal pausing and RNAPII processivity, mechanisms that are hard to measure through bulk RNA analysis. We observed that B02-treated cells showed lower overall levels of nascent transcription peaks on the viral genome (**Figure 3D**). We quantified the total number of peaks specifically on IE and E genes (**Figure 3E**) and found that B02 treatment caused a decrease in nascent transcription of IE genes, suggesting its role in initiation of viral transcription. Similar effects were also observed in HFFs (data not shown). To probe this further, we compared the PRO-seq peaks on gene promoters versus gene bodies and observed a higher impact of RAD51 inhibition on promoters compared to gene bodies (**Figure 3F)**. In summary, these data suggest that RAD51 mediates viral transcription initiation, through its interactions on IE gene promoters.

**Figure 3:**
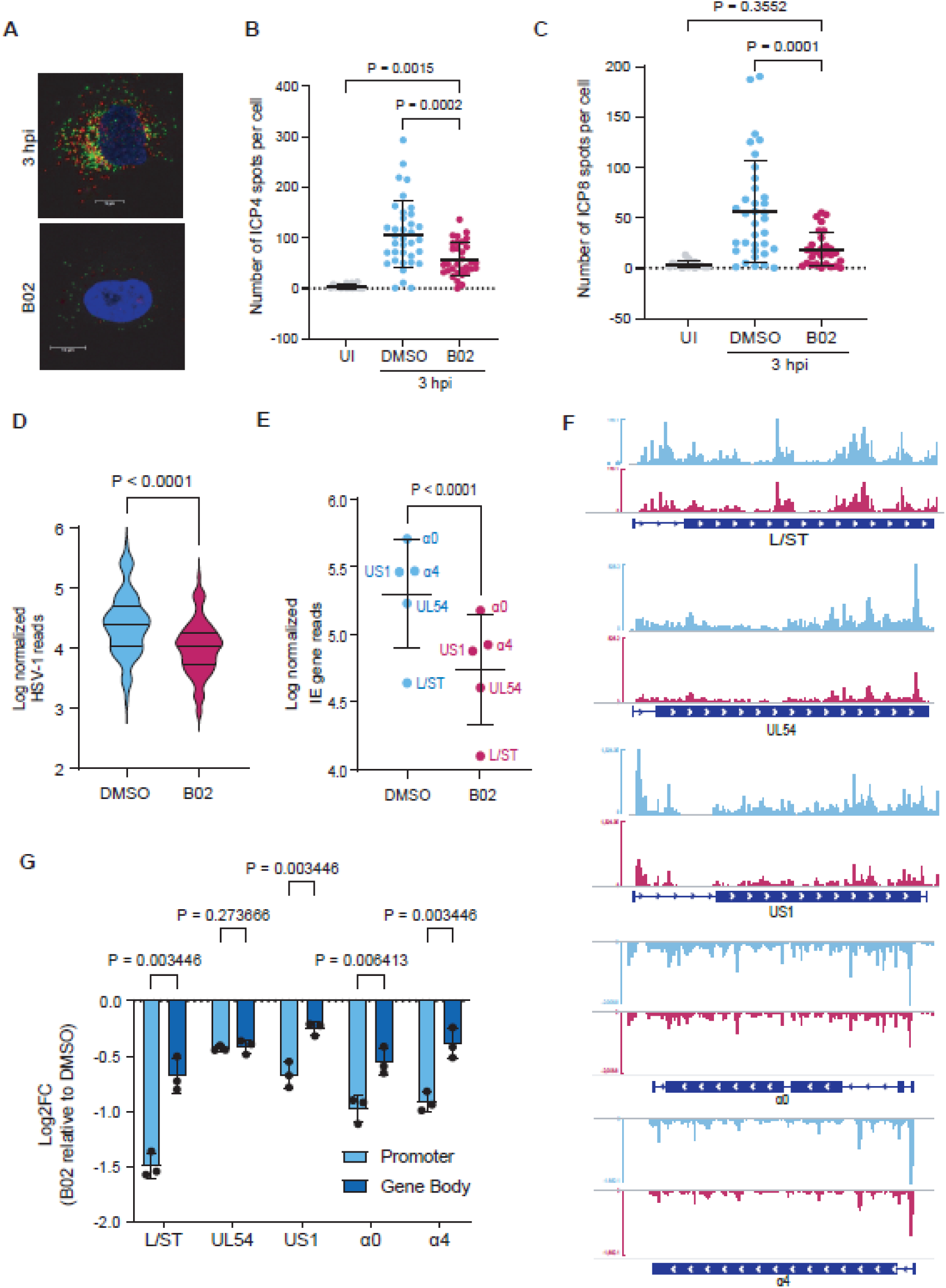
RAD51 inhibition leads to decreased nascent viral transcription. **A**. HFF cells were treated with DMSO or 20uM B02 for 2 hours pre-infection. Cells were then infected with HSV-1 17syn+ at MOI 5 and fixed for single-molecule RNA FISH at 3 hpi. **B-C**. ICP4 and ICP8 foci per cell were quantified. Statistics: One-Way-ANOVA **D.** HEp2 cells were pre-treated with DMSO or B02 for 16 hours. Cells were then infected with HSV-1 strain F at MOI 5 and harvested at 1.5 hpi for PRO-seq analysis. Plot shows the total log normalized RNAP II reads in DMSO or B02 treated cells. Statistics: Student’s T-test. **E**. Quantification of log normalized immediate-early gene reads in DMSO or B02 treated cells**. F.** PRO-seq peaks for immediate-early genes. Statistics: Paired Student’s t-test G. Fold-change (B02/ DMSO) of RNAP II reads on immediate-early genes at gene promoter vs gene body. Statistics: Two-Way-ANOVA.

### RAD51 interacts with host and viral transcription factors to promote viral transcription

The HSV-1 IE gene promoters are known to be GC-rich and serve as landing sites for viral and host transcription factors (59,60). Upon nuclear entry of the viral dsDNA, HSV-1 IE gene expression is mediated by the viral tegument protein VP16 and host transcription factors (Oct-1, Sp1, HCF-1) that assemble as a complex and limits the assembly of heterochromatin on genomes (32,59,61). We performed an Immunoprecipitation followed by immunoblot analysis to examine the interaction of RAD51 protein with IE viral transcription factors VP16 and ICP4. We found that RAD51 interacted with VP16 and ICP4 and that this interaction was decreased by B02 treatment (**Supplementary Figure S3A)** Interaction with ICP4 was also significantly decreased when viral transcription was inhibited using 5,6-dichloro-1-β-D-ribofuranosylbenzimidazole (DRB). However, RAD51 interacted with VP16 even in the absence of active viral transcription, suggesting they interact on incoming genomes. In contrast, no interaction was observed with the virus-encoded ubiquitin E3 ligase ICP0, suggesting RAD51 interactions are specific to viral transcription. To identify regulatory factors associated with RAD51 at gene promoters, we performed a motif analysis on the PRO-seq peaks on the IE promoters that were most affected by RAD51 inhibition. GC-rich motifs resembling Sp1/KLF family binding sites were highly enriched, suggesting that RAD51 might bind GC-rich structures at promoter regions (**Figure 4A**). We cannot exclude the possibility that this enrichment reflects underlying sequence composition rather than specific transcription factor recruitment. However, consistent with our motif analysis, RAD51 and BRCA2 have previously been shown to interact with Sp1 transcription factor (62). GC-rich regions also have the potential to form secondary structures such as G-quadruplexes (G4s). It has been previously shown that HSV-1 genome contain G4s and acts as landing sites for ICP4 to mediate transcription (60). To examine whether RAD51 acts on G4 forming regions, we first blocked access to G4s using G4 stabilizers, Pyridostatin (PDS) and N-methyl mesoporphyrin IX (NMM) (**Supplementary Figure S3B**). As previously shown, G4 stabilizers led to a decrease in viral progeny production (26), however when combined with RAD51 knockdown, we did not observe an additive effect, suggesting that RAD51 may be functioning at these G-rich secondary structures. However, these data do not provide direct evidence for RAD51 binding G4 structures. To test further the ability of RAD51 to promote gene expression directly, we cloned the ICP4 enhancer region (ICP4e) containing Sp1, VP16 and ICP4 binding sites into a plasmid containing a luciferase reporter gene (**Figure 4B**). We showed that while RAD51 inhibition led to a 2-fold decrease in luciferase expression (**Figure 4C**), overexpression of RAD51 led to a dose-dependent increase in luciferase expression (**Figure 4D**). To examine direct binding of RAD51 to the ICP4e region, we performed CUT&RUN-qPCR using the WT or ICP4e regions with Sp1, VP16 or ICP4 sites mutated **(Figure 4E).** We found that RAD51 bound ICP4e regions, and that mutating the Sp1 or VP16 binding sites partially abrogated the binding, while mutating the ICP4 binding region had a milder effect (**Figure 4F)**. Finally, we expressed the WT and Sp1 mutant ICP4e constructs and measured luciferase expression when RAD51 was inhibited. While Sp1 binding mutant and RAD51 inhibition (B02) displayed a similar defect in luciferase expression, the combination of Sp1 binding site mutation and B02 treatment was not synergistic, indicating that RAD51 might act as a complex with viral and host transcription factors (**Figure 4G**). Together, these data suggest that RAD51 cooperates with viral and host transcription factors VP16 and Sp1/KLF4 to drive IE gene expression from incoming viral genomes.

**Figure 4:**
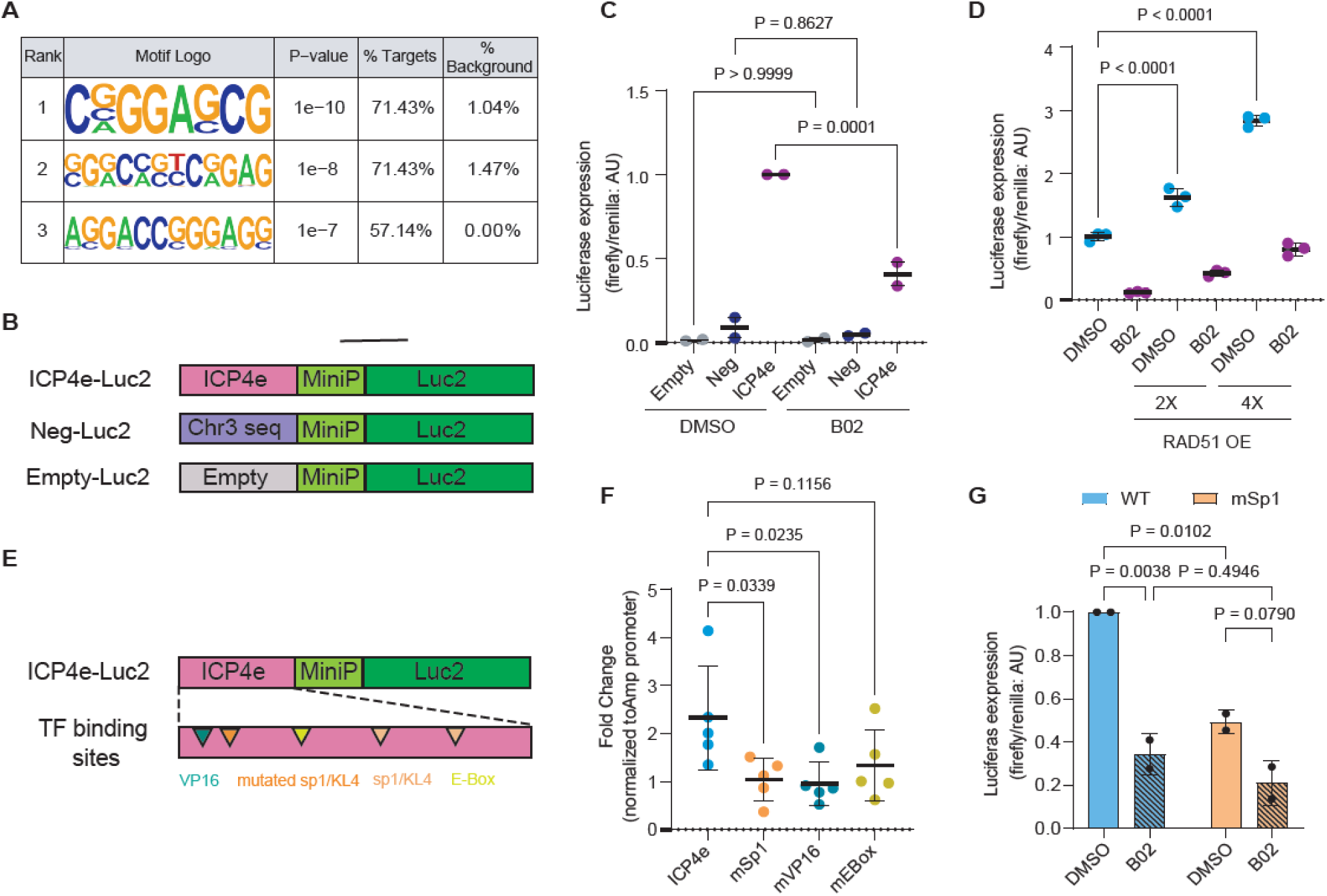
RAD51 interacts with early host and viral transcription factors to promote viral transcription. **A**. Motif analysis performed on IE gene promoters that were changed in the PRO-seq analysis when RAD51 was inhibited. **B**. Luciferase plasmid constructs used to analyze the effect of RAD51 on gene expression. **C**. HEp2 cells were transfected with indicated Luciferase plasmids. 24 hours later, luciferase expression was read to examine the impact of RAD51 inhibition on gene expression. Cells were pre-treated with DMSO or B02 for 16 hours before the assay. Statistics: One-Way-ANOVA. **D**. HEp2 cells were co-transfected with indicated ICP4e luciferase plasmid and a RAD51 overexpression plasmid. 24 hours later, luciferase expression was read to examine the impact of RAD51 overexpression on gene expression. Statistics: One-Way-ANOVA. **E**. Mutant Luciferase plasmid constructs used to assess gene expression and RAD51 binding. **F**. HEp2 cells were transfected with indicated plasmids mutated for sp1, VP16 or ICP4 (Ebox) binding sites. RAD51 binding to the ICP4e region was assessed using CUT&RUN-qPCR. Statistics: One-Way-ANOVA. **G**. HEp2 cells were transfected with the WT ICP4e or sp1 mutated ICP4e plasmid. Cells were pre-treated with DMSO or B02 for 16 hours and luciferase expression was measured 24 hours post transfection. Statistics: Two-Way-ANOVA.

### RAD51 interactome reveals interaction with ADNP, a core factor of the ChAHP complex

To identify other host proteins associated with RAD51 during HSV-1 infection, we performed RAD51 immunoprecipitation followed by mass spectrometry (IP-MS) from infected cells. We identified known RAD51 interactors, BRCA2 and PALB2 in our dataset (63). We also identified a previously unknown significant interaction with ADNP and CBX3 (HP1gamma) in uninfected as well as HSV-1 infected cells (**Figure 5a, Supplementary Fig S3C, S3D)**. Specifically, we observed a 2-fold increase in ADNP interaction during infection compared to uninfected cells (data not shown). ADNP is a core protein of the ChAHP complex, consisting of CHD4, ADNP and HP1 proteins, that is known to regulate transcription by repressing transcription through recruitment of HP1 and CHD4 to specific genomic sites (64,65). To assess whether the interaction is functionally relevant during infection, we performed a double knockdown of RAD51 and ADNP and measured viral progeny and protein production **(Figure 5B, 5C).** While ADNP knockdown alone did not have a significant effect on viral protein levels and progeny production in control cells, when knocked down in RAD51-deficient cells, ADNP was able to completely rescue viral protein levels and partially rescue infectious progeny production (**Figure 5B, 5C**). These data indicate that RAD51 acts to suppress the repressive function of ADNP and potentially the ChAHP complex on incoming genomes, promoting access to viral and host transcription factors to drive IE gene expression.

**Figure 5:**
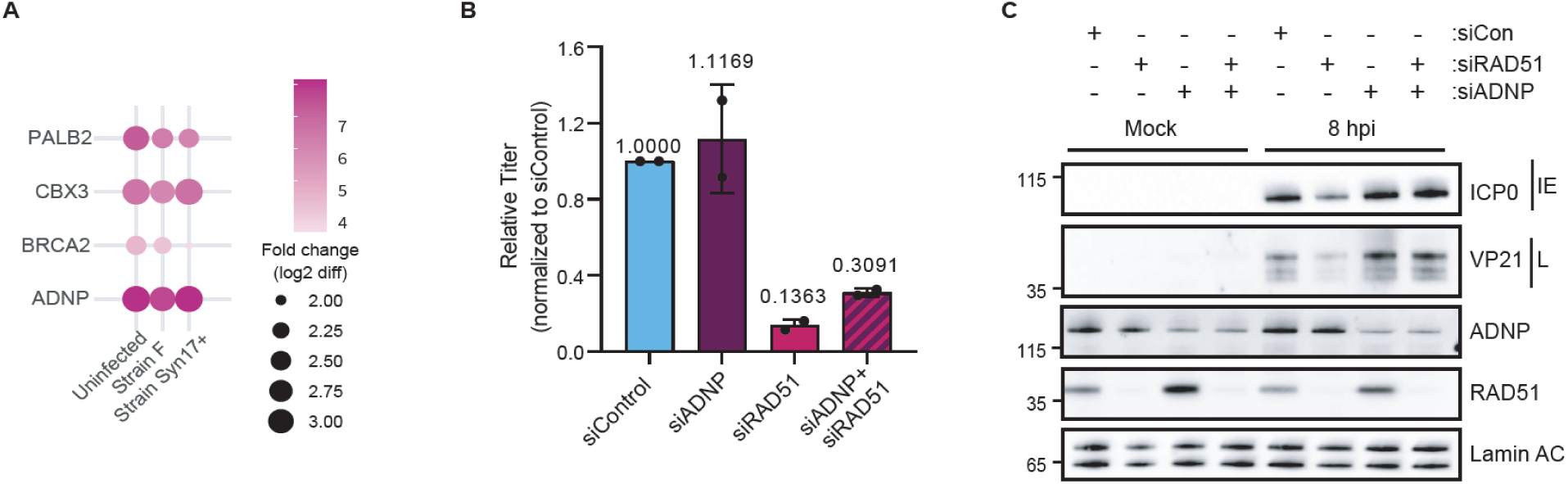
RAD51 interacts with the ADNP: a core factor of the ChAHP complex. **A**. HEp2 cells were treated with DMSO/ B02 for 16 hours or infected with HSV-1 strain 17syn+ for 3 hours and strain F for 1.5 hours. Cells were harvested for RAD51 Immunoprecipitation followed by mass spectrometry (IP-MS). Dot plot showing fold-change over IgG of RAD51 interactors identified in uninfected and HSV-1 infected cells. BRCA2 and PALB2 are known RAD51 interactors. ADNP and CBX3 are part of the ChAHP complex. **B**. HFF cells were transfected with control, ADNP, RAD51 or ADNP+RAD51 siRNA. 48 hours post transfection, cells were infected with HSV-1 17syn+ at MOI 0.1 and harvested for plaque assay at 24 hpi. Plaque assay was performed in Vero cells and plaques were counted after 3-4 days. **C.** HFF cells were transfected with control, ADNP, RAD51 or ADNP+RAD51 siRNA. 48 hours post transfection, cells were infected with HSV-1 17syn+ at MOI 3 and harvested for western blot at 8 hpi.

## Discussion

RAD51 has well established roles during HR and at DNA replication forks (16). Here we reveal a previously unknown function of RAD51 in promoting transcription of HSV-1 gene expression. We show that RAD51 inhibition or deficiency leads to overall lower viral progeny and protein production, which is distinct from its known roles in DNA recombination and replication. We observed that RAD51 inhibition led to decreased IE gene expression, with a more prominent effect on gene promoters, highlighting its role in transcription initiation. We also observed RAD51 binding to the ICP4 gene regulatory region, which was diminished when early transcription factor (TF) sites were mutated, suggesting a potential complex with TFs at these regions. Finally, we uncovered a previously unknown interaction of RAD51 with ADNP and CBX3, part of the ChAHP transcription repression complex. We propose that RAD51 binding to IE viral gene promoters inhibit recruitment of the ChAHP complex, thus allowing for transcription factor access to the genomes (**Figure 6**). However, there are some outstanding questions remaining to be addressed: **1)** The direct evidence for RAD51 binding to a specific DNA substrate or structure on the viral genome is lacking. **2)** It is not known the extent to which RAD51 and ADNP compete for the same DNA substrate or if their interaction is independent of DNA binding. **3)** There is very little evidence for chromatinization of viral genomes during lytic infection (66). Considering the association with ADNP, it would be worth examining the viral chromatin accessibility when RAD51 is inhibited.

**Figure 6:**
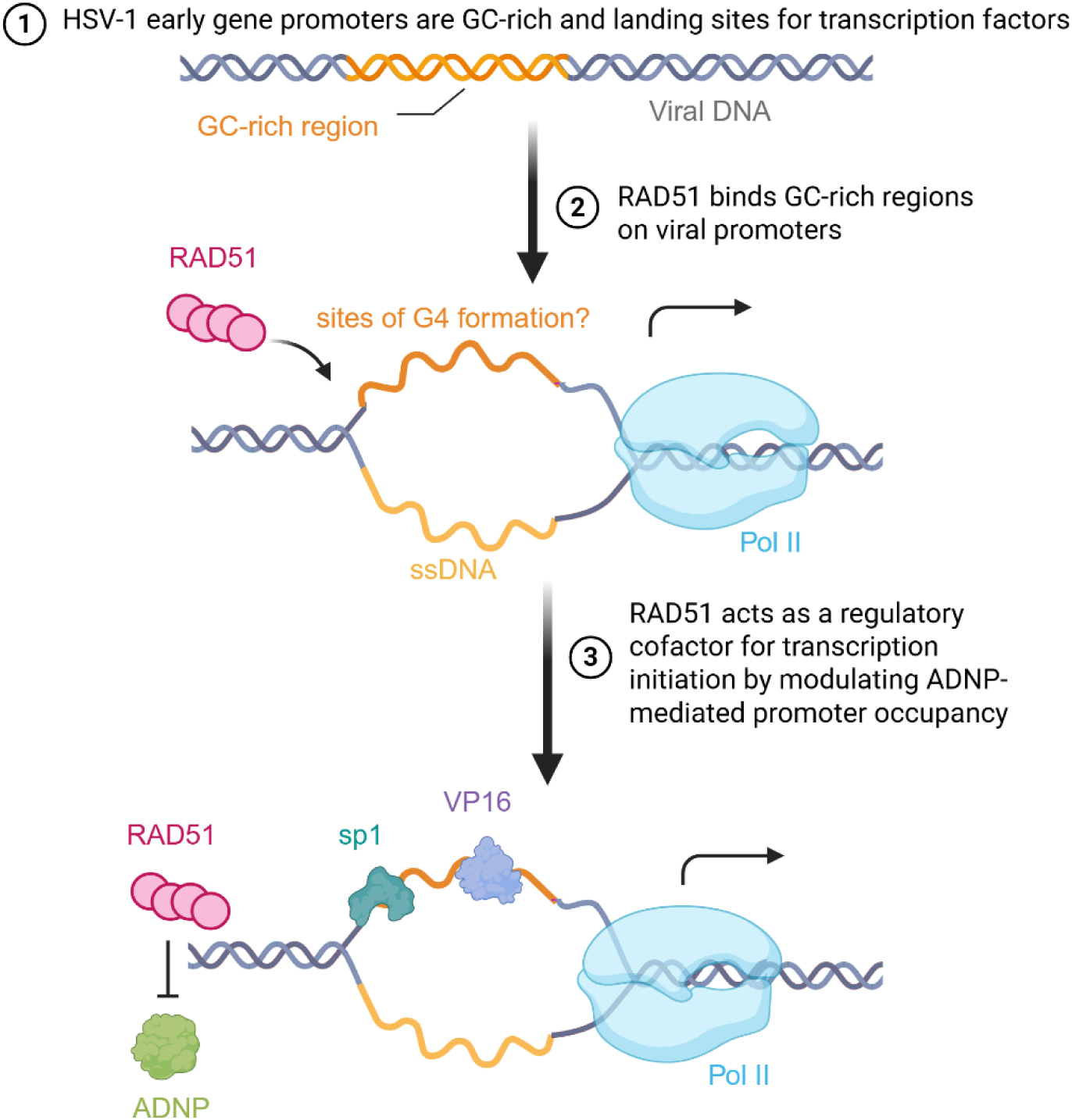
**Working model highlighting the role of RAD51 during HSV-1 infection in promoting early HSV-1 transcription.**

Overall, our findings reveal a new mechanism for viral transcriptional regulation and highlight how virus infection hijacks cellular machinery for its benefit. We have revealed a non-canonical role for RAD51, independent of its role in DNA repair and replication. We hypothesize that RAD51 allows for viral chromatin accessibility by recognizing and directly binding DNA at GC-rich viral promoters. Moreover, this role of RAD51 in driving transcription has important implications for the fields of genome instability and cancer and should therefore be explored outside the context of virus infection.

### DNA repair proteins are manipulated during early stages of infection

Incoming HSV-1 genomes were shown to contain nicks and gaps, which are suggested to recruit DNA repair proteins during very early stages of infection (67). However, when these genomes were repaired prior to infection, they did not have any impact on infectivity (67). Therefore, it is still not clear whether these proteins are recruited to repair damage on incoming genomes, or whether they perform other vital functions necessary for productive infection. In addition, HSV-1 induces a DDR response that is distinct from conventional DDR response induced by a DNA damaging agent (47,48). We also show that while RAD51-deficient cells produce more than 10-fold fewer progeny, knockdown of RPA, does not result in a similar decrease of infectious progeny. These studies indicate that the DDR response during HSV-1 infection is complex and the role of individual factors must be systematically elucidated.

HSV-1 genomes are GC-rich and contain repeat sequences, specifically around IE gene regions. GC-rich regions, specifically containing G-runs, are prone to form secondary structures such as G-quadruplexes (27,60,68). G-quadplexes can act as landing sites for transcription factors and regulate gene expression (69). Similarly, during HSV-1 infection, ICP4 has been shown to bind and unfold G-quadruplexes to allow for transcription (60). Here we show the recruitment of RAD51 to GC-rich regions on viral promoters and demonstrate a direct role in activating gene expression. We propose that RAD51 binds secondary structures on GC-rich promoters and helps recruit early viral and host transcription factors. RAD51 is an ATPase with high affinity to ssDNA (15,17). It is possible that RAD51 binds ssDNA DNA at regions of G4 formation and facilitates unwinding through its ATPase activity. We propose that virus infection is an attractive tractable system to reveal new roles for DDR proteins at DNA secondary structures.

### Expanding roles for DNA repair proteins in transcription

There is a strong interplay between transcription and DNA damage and repair (70). Studies over decades have demonstrated that the presence of DNA damage stalls RNAPII and promotes recruitment of DNA repair proteins (71,72). We have shown, on the viral genome, that inhibition of RAD51 did not increase RNAPII pausing, suggesting that it is not unrepaired DNA damage which leads to decreased viral transcription. Supporting this, we also observed no change in viral DNA recombination when RAD51 was inhibited. In contrast to our findings, it was shown that RAD51 can promote oncogene expression through its function at DSBs on promoter enhancer regions (73). Together, these data point to a major gap in our understanding of how DNA repair proteins cooperate with transcription and how these two processes are more intertwined than previously understood.

Recent studies have focused on roles for DNA repair proteins in directly promoting transcription, acting as transcription factors or cofactors and modulating the chromatin structure (70). Factors from the homologous recombination (HR), nucleotide excision repair (NER), Fanconi Anemia (FA), and Single-Stranded Break (SSB) repair pathways have all independently been shown to promote transcription. Transcriptional activation involves the initial recognition of key regulatory DNA elements at promoters by sequence or structure-specific DNA-binding activators and the core transcription machinery, along with the recruitment of essential cofactors. The role of regulatory structures at transcription sites is being studied only recently. It is highly possible that DNA repair proteins play a crucial role in recognizing these secondary structures at gene regulatory elements and elicit a chromatin environment that facilitates transcription initiation. In support of this hypothesis, base excision repair (BER) proteins were shown to regulate G4 formation and G4-mediated gene expression (74).

### Role for DNA repair proteins in reactivation from latency

HSV-1 can establish latency in neurons(75,76). Latent genomes are chromatinized, suppressing viral gene expression except for except for Latency Associated Transcripts (LATs) (76). Reactivation requires removal of heterochromatin and expression of IE genes (66,77,78). We show that RAD51 inhibition impacts transcription initiation at viral IE gene promoters, potentially preventing repression by the ChAHP complex. It would be interesting to investigate the role of RAD51 during latency and whether it impacts either maintenance of latency or reactivation from latency. Additionally, the role of the ChAHP complex in promoting latency would be an interesting to address in future studies. Broadly, DDR in post mitotic neurons is not completely understood (79), and more research is required to understand the interactions of DDR proteins with viral genomes during latency.

### Limitations of the study

Our study reveals a DNA damage-independent role for RAD51 at viral gene promoters. We show that RAD51 inhibition led to decreased nascent transcription. RAD51 binds GC-rich regions and drives active transcription. We propose that RAD51 binds secondary structures such as G-quadruplexes at promoters and acts as transcription cofactor by allowing transcription factor complex assembly. However, this model still needs to be tested fully as there is no direct evidence yet for RAD51 recognizing viral G-quadruplexes. We also employed IP-MS to identify RAD51 interactors. While we saw positive hits BRCA2 and PALB2, and identified a novel interaction with ADNP, our overall dataset was limited due to poor antibody binding. Future studies will focus on generating better tools and reagents to study RAD51 interactome to gain a deeper understanding of its non-canonical roles.

## Acknowledgements

We would like to acknowledge Dr Roger Greenberg for providing us with PEO1 WT and BRCA KO cells. We are grateful to Dr. Kavitha Sarma for discussions regarding ADNP. We thank Emily Alchaer for assistance with western blots and immunofluorescence. We thank Cassandra Wong of the Network Biology Collaborative Centre Proteomics Facility (RRID: SCR_025375) at the Lunenfeld-Tanenbaum Research Institute for mass spectrometry analysis. The facility is supported by the Canada Foundation for Innovation and the Ontario Government. We thank members of the Weitzman lab for helpful feedback and discussions.

## Contributions

N.K conceptualized and designed the study with feedback from M.D.W. M.D.W provided funding and resources. L.E.M.D performed the PRO-seq. JDB supervised and provided resources for PRO-seq. L.E.M.D, K.E.H, and N.K performed the PRO-seq analysis. T.M.T performed and analyzed the RAD51 IP-MS. M.R.A, S.S, designed, performed and analyzed the smRNA FISH and Recombinant plaque assay with supervision from OK. Y.A developed software for recombinant plaque analysis and J.K modified software for smRNA FISH analysis. B.G.W performed the RAD51 CUT&RUN-qPCR. H.M.C and E.H performed western blots. N.K performed all other experiments. N.K wrote the manuscript draft with feedback from M.D.W. All authors read and provided feedback on the manuscript.

## Materials and Methods

### Cell culture

Primary human foreskin fibroblasts (HFFs), BJ, MRC5, Hep2, Vero were obtained from the American Type Culture Collection (ATCC). Cells were grown DMEM (Gibco) supplemented with 10% fetal bovine serum (FBS) (VWR) and penicillin (100 U/ml)/streptomycin (100 μg/ml) (Invitrogen) in a 5% CO2 humidified incubator at 37°C.

### siRNA and plasmid transfections

Gene knockdown experiments by siRNA were carried out using Lipofectamine RNAiMAX transfection reagent (Invitrogen), following the manufacturer’s instructions. siRNA for siGENOME non-targeting control, RAD51, RPA, and ADNP were purchased from Dharmacon/ Horizon Discovery. Plasmid transfections were performed using Lipofectamine 2000 transfection reagent, according to the manufacturer’s instructions. The pGL4.23[luc2/minP] Vector was purchased from Promega.

### Viruses and Infection

The HSV-1 strain 17syn+ and strain F were propagated and titrated in Vero cells. Recombinant viral constructs expressing fluorescent proteins are derivatives of HSV-1 strain 17syn+. OK11 contains mCherry fluorescent protein with a nuclear localization tag under the CMV promoter between UL37 and UL38 genes. OK42 expresses both mTurq2 (CFP) and mVenus (YFP) genes located between UL37 and UL38 genes and fused within the UL25 gene, respectively (57). Monolayers of cells were infected with HSV-1 at an indicated MOIs in DMEM supplemented with 2% FBS. Cells were incubated at 37 °C for 1 h to allow for virus adsorption. After 1 h, the inoculum was removed and replaced with DMEM with 10% serum, and infection was allowed to proceed. This was considered 1 hour post infection (hpi).

### Plaque assay

Monolayers of cells were infected with HSV-1 at indicated MOI 0.1 DMEM supplemented with 2% FBS. Cells were incubated at 37 °C for 1 h to allow for virus adsorption. After 1 h, the inoculum was removed and replaced with DMEM with 10% serum, and infection was allowed to proceed. Infected cells were harvested at 24 hpi by scraping into media, and viruses were released from cells by 3Xfreeze-thaw in LN_2_ and 37°C water bath. Cells were centrifuged at 10,000 RPM for 10 minutes to collect cell debris and supernatants were diluted serially to infect Vero cells. After 1 hour, cells were overlaid with medium containing 0.5% carboxylmethylcellulose. Plaques were stained with crystal violet at 3 days post infection.

### Immunoblot analysis

Cells were washed with PBS and scraped using NuPAGE 1X LDS Sample Buffer (Invitrogen). cell extracts were prepared by boiling the samples for 10 minutes at 95°C. Proteins were separated via SDS-PAGE and visualized using SuperSignal WestPico Chemiluminescent Substrate (Thermo Scientific) and GBox imaging system (Syngene).

### Immunofluorescence

Cells were plated on glass coverslips in 24-well plates and infected with HSV-1 MOI 3. Cells were washed in PBS, fixed in 4% paraformaldehyde for 10 min, permeabilized with 0.5% Triton X-100 in PBS for 10 min. Cells were pre-extracted prior to fixation for RAD51 immunofluorescence using CSK buffer (100 mM NaCl, 300 mM sucrose, 10 mM PIPES, pH 6.8, 3mM MgCl2, and 0.5% Triton X 100). Cells were blocked with 3% bovine serum albumin (BSA) with 5% goat serum for 1h. Cells were incubated with primary antibodies overnight at 4C. Next day, Alexa Fluor-conjugated secondary antibodies were incubated for an hour at room temperature. Nuclei were visualized by staining with 4’,6-diamidino-2-phenylindol (DAPI). Coverslips were mounted using ProLong Glass Antifade Reagent and immunofluorescence was visualized using a Zeiss LSM 710 or LSM880 Confocal microscope (Cell and Developmental Microscopy Core at UPenn) and ZEN software. Images were processed using ImageJ.

### Single-molecule RNA FISH

HFF cells were grown in 12-well slide chambers and were incubated in the presence or absence of 20µM B02 for 2 hours pre-infection, then infected with HSV-1 at MOI 5. Cells were fixed with 4% PFA for 10 min at 3hpi. smRNA FISH was carried out for ICP4 and ICP8 using ViewRNA ISH Cell Assay Kit by Invitrogen (ThermoFisher QVC001) according to the manufacturer’s instructions. Images were taken using SP8 Leica Confocal Microscope and were analyzed by modified quant FISH/ big FISH using Python.

### Recombinant plaque assay

HFF cells were plated in 12-well plates. Two hours pre-infection, media was incubated with DMSO or 20uM RAD51 inhibitor B02. Cells were co-infected with an equal mix of OK11 and OK42 at MOI 20 and incubated at 37 °C for 1 hour. Following incubation, inoculum was removed, washed three times with PBS and fresh media was added, with or without inhibitors. After incubation for 18 hours at 37 °C, the media was collected and stored at −80 °C. Samples were sonicated and centrifuged to pellet any cell debris and the supernatant was diluted to infect confluent Vero cells in 6-well plates. 36-48 hours post infection, 4 regions were imaged from each well. Viral Plaques were imaged using Nikon Eclipse Ti-E epifluorescence inverted microscope (Nikon, Tokyo, Japan). Plaquer analysis was performed as described in (57).

### PRO-seq

HEp2 or HFF cells were incubated with DMSO or B02 for 16 hours pre-infection. Cells were infected with HFF strain F (HEp2) or strain 17syn+ (HFF) at MOI 5 and harvested at 1.5 hpi or 3 hpi respectively. Nuclei were isolated from infected cells as previously described (80,81). Cells were washed 2x with ice-cold PBS and incubated for 10 min on ice with swelling buffer (10 mM Tris-HCl [pH 7.5], 10% glycerol, 3 mM CaCl2, 2 mM, MgCl2, 0.5 mM DTT, protease inhibitors [Pierce] and 4 U/mL RNase inhibitors). Cells were scraped from the plate, pelleted via centrifugation at 600× g for 10 min (4 °C), resuspended in lysis buffer (swelling buffer + 0.5% Igepal), and then incubated on ice for 20 min to release nuclei. Nuclei were pelleted by centrifugation at 1500× g for 5 min (4 °C), washed 2x with lysis buffer with a final wash in storage buffer (50 mM Tris-HCl [pH 8.0], 25% glycerol, 5 mM MgAcetate, 0.1 mM EDTA, 5 mM DTT). Nuclear run-on and PRO-seq library preparation was performed as described previously (82). For analysis, FASTQ files were processed using the PRO-Seq 2.0 pipeline, developed by the Danko lab at Cornell: https://github.com/Danko-Lab/proseq2.0. The genome used to align reads was a concatenated file containing hg38 and HSV-1 F or HSV-1 17syn+ genomes. The external repeats on the viral genome were deleted to aid sequencing alignment. Data were normalized for sequencing depth based on total paired reads and viral genome copy number. HSV-1 normalized bigwig files were visualized using IGV genome browser.

### Co-immunoprecipitation (Co-IP)

8 million Hep2 cells were infected with HSV-1 at MOI 10. Cells were treated with DMSO or B02 16 hours pre-treatment. 5,6-dichloro-1-β-D-ribofuranosylbenzimidazole (DRB) was added 1 hour post infection. Cells were harvested in 500μl of ice-cold co-IP buffer (50 mM Tris-HCl, pH 7.4, 150 mM NaCl, 1 mM EDTA, 10% glycerol, 0.1% NP40/IGEPAL). Protease inhibitors were added at 1X final concentration. Cell pellets were frozen on dry ice (∼10min) and thawed in warm water bath (37C) with constant shaking, and transferred to ice. Samples were rotated end over end for 15-20min at 4C. The 20 μl of Protein G Dynabeads were incubated with RAD51 antibody for 1 hour at room temperature. Samples were cleared by centrifugation and incubated with beads overnight at 4°C with constant rotation. The beads were then washed with co-IP buffer (lysis buffer without inhibitors) and resuspended in 1X LSD sample buffer.

### Luciferase assays

The ICP4e WT and mutant regions were synthesized by Twist Biosciences and cloned into the KpnI and BglII sites of pGL4.23[luc2/minP]. HEp2 cells were co-transfected with pGL4.23[luc2/minP] and a pRL *Renilla* Luciferase Control Reporter plasmid in a 96-well plate. Cells were teated with DMSO or B02 for 16 hours pre-infection. 24 hours post transfection, luciferase assay was performed using the Dual-Luciferase® Reporter Assay System, Firefly, Renilla (Promega), according to the manufacturer’s instructions. The Renilla expression was used for normalization.

### CUT&RUN-qPCR

HEp2 cells were transfected with 2ug of the pGL4.23[luc2/minP] plasmid containing the ICP4e WT or mutants. Next day, cells were harvested for CUT&RUN using the Epicypher CUTANA™ ChIC/CUT&RUN Kit, according to the manufacturer’s protocol. This was followed by qPCR to amplify the ICP4e region using Luc2 primers. Samples were normalized to IgG and to the Ampicillin promoter on the plasmid to normalize to plasmid copy number.

Luc2 FP: TCTGGCCTAACTGGCCGGTA

Luc2 RP: ACAGTACCGGATTGCCAAG

Amp Promoter FP: CTTACTTATCCTTGAGAGACGTACTAG

Amp Promoter RP: CAACCAAGTCGTTTTGTGAGTAGTG

### RAD51 Immunoprecipitation Mass Spectrometry

Forty million HEp2 cells were either treated with B02 or DMSO 16 hours pre-infection or infected with HSV-1 17syn+ for 3 hours or strain F for 1.5 hours. Prior to immunoprecipitation nuclei were enriched following the REAP method (21067583), by briefly washing cells in 0.1% NP-40 in PBS supplemented with protease inhibitor cocktail. Nuclei were immediately resuspended in NP-40 lysis buffer and snap frozen on dry ice. Samples were then partially thawed in a 37°C water bath and then incubated at 4°C for 20 minutes with end over end rotation. Lysed cells were centrifuged to clear cellular debris and supernatant was transferred to 1.5mL Lobind Eppendorf tubes. Protein G Dynabead-antibody complexes were prepared by mixing 1.5 ug RAD51 antibody (Sigma; PC130) or IgG control (Cell Signaling Technology; 3900S) with 15 uL Protein G dynabead in NP-40 lysis buffer for 1 hour at room temperature. Following incubation, beads were washed in NP-40 lysis buffer and resuspended with lysate. Immunoprecipitations were carried out at 4°C for 4 hours with end over end rotation. Immunoprecipitated complexes were washed twice in NP-40 lysis buffer and then an additional two times in Rinse Buffer. Beads were then resuspended in 100 uL of trypsin digest buffer containing 1 ug trypsin (Promega; V5111) and incubated at 37°C overnight on a thermomixer. The following day 0.5 ug trypsin was added and incubated for an additional 3 hours. Supernatants containing digested peptides were transferred to new Lobind Eppendorf tubes, acidified with formic acid to a 5% final concentration, and dried in a vacuum centrifuge.

NP-40 lysis: 50mM Tris-HCl pH 7.4, 150mM NaCl, 1mM EDTA, 10% glycerol, 0.1% NP-40, 25U benzonase/mL (1:1000 of 250K unit tube), 1x HALT protease/phosphatase inhibitor

Rinse Buffer: 20mM Tris-HCl pH 8, 2mM CaCl2

Digest buffer: 20mM Tris-HCl pH 8

### Mass spectrometry and Analysis

For data-dependent acquisition (DDA) LC-MS/MS, one-twenty fourth of digested peptides were analyzed using a nano-HPLC (High-performance liquid chromatography) coupled to MS. Sample was loaded onto Evotip Pure per manufacturer instructions. Peptides were eluted from the Performance Column (cat#: EV-1109, 8cmx150µm with 1.5µm beads, heated at 40C) with the 60SPD pre-formed acetonitrile gradient generated by an Evosep One system, and analyzed on a timsTOF Pro 2. The Evosep was coupled to timsTOF Pro 2 using a 20um diameter emitter tip. The column toaster was set to 40C. The total DDA protocol is 22 minutes. The MS1 scan had a mass range of 100-1700Da in PASEF mode. TIMS settings were accumulation and ramp time of 100ms (with 4 PASEF ramps and active exclusion at 0.4min), and within the mobility range (1/K0) of 0.85 to 1.3V·s/cm2. This was at a cycle time of 0.53s. The target intensity was set to 17,500 and intensity threshold set to 1750. 1+ ions are excluded from fragmentation using a polygonal filter. The auto calibration was off. MSFragger 4.1 was used to search mass spectra using a FASTA database consisting of human (UP000005640) and human herpesvirus 1 (UP000180758) proteins, no isoforms. Acetylated protein N-term and oxidated methionine were set as a variable modification. Precursor mass tolerance was set to 20ppm on either side. Fragment mass tolerance was set to 20ppm. Enzymatic cleavage was set to trypsin with 2 missed cleavages. MSBooster and Percolator were turned on. Percolator required a minimum probability of 0.5 and did not remove redundant peptides. The target-decoy competition method was used to assign q-values and PEPs. For ProteinProphet, the maximum peptide mass difference was set to 30ppm. When generating the final report, the protein FDR filter was set to 0.01. FDR was estimated by using both filtered PSM and protein lists. Razor peptides were used for protein FDR scoring. All other parameters were default. All other parameters were default.

Protein groups with high missing values (>50%) and contaminants were filtered prior to downstream analysis. Protein intensities were transformed (log2), vsn normalized, and imputed using the minimum probability distribution. Differential protein abundance was performed with Limma (ref) to determine high-confidence protein interactors compared to the IgG control. Proteins with a log2 fold-change ≥ 1 and p value ≤ 0.05 were considered to be high-confidence.

### Statistical Analysis

Graphs and statistical analysis were performed using GraphPad Prism 11.0.2. The statistical analysis of Plaquer performance was done with Python (version 3.11) and SciPy (version 1.13.1).

## Supplementary Figure Legends

**Figure S1:**
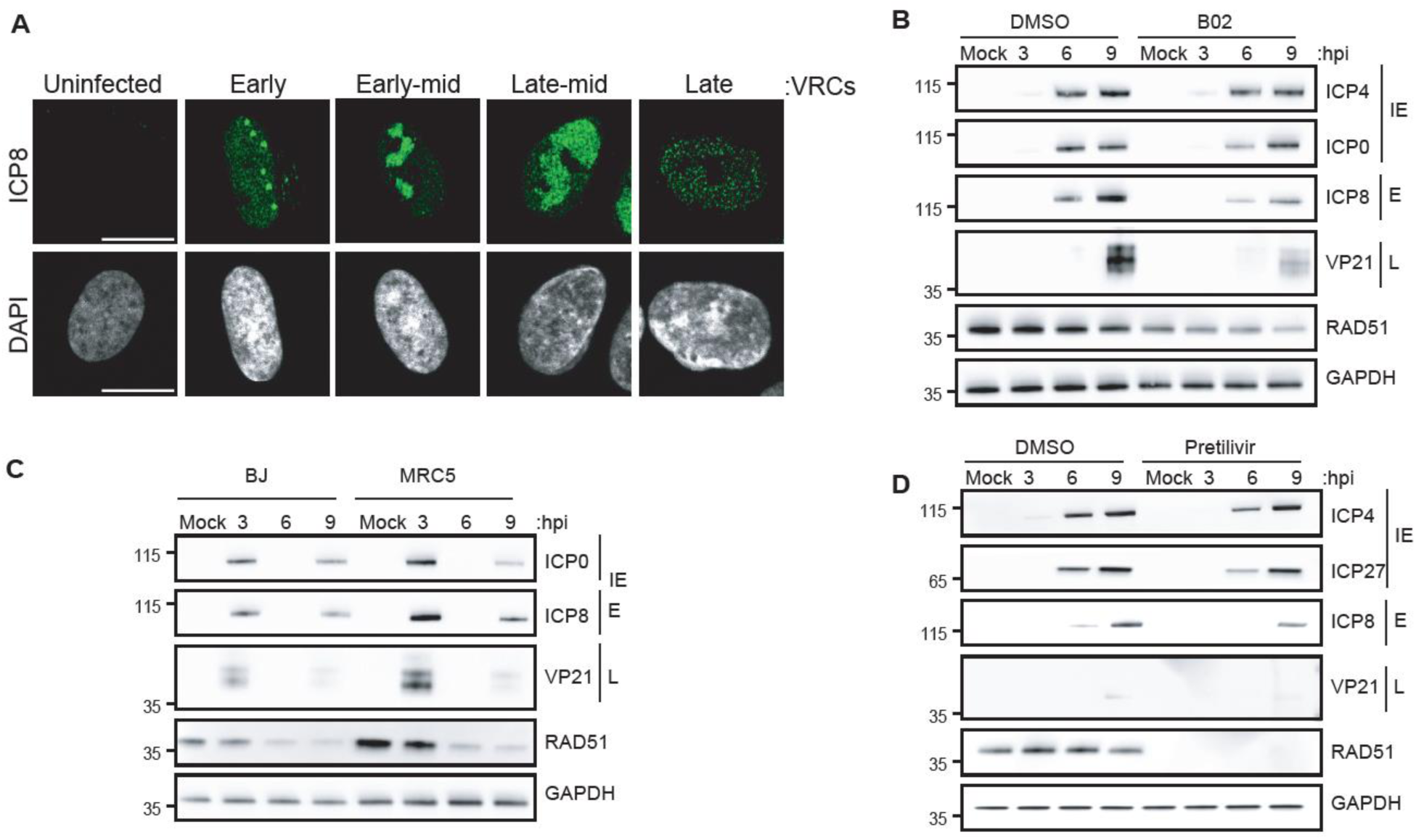
RAD51 loss leads to decreased HSV-1 progeny and protein production. **A.** Cells representing different stages of viral replication compartments (VRCs). **B.** HFF cells were pre-treated with DMSO or B02 for 16 hours. Cells were then infected with HSV-1 17syn+ at MOI 3 and harvested for western blot at 3, 6, 9 hpi. **B.** BJ and MRC5 cells were transfected with a control or RAD51 siRNA. 48 hours post transfection, cells were infected with HSV-1 17syn+ at MOI 3 and harvested for western blot at 3, 6, 9 hpi. **C.** HFF cells were transfected with a control or RAD51 siRNA. 48 hours post transfection, cells were infected with HSV-1 17syn+ at MOI 3 and harvested for western blot at 3, 6, 9 hpi. Pretilivir was added at 1 hpi.

**Figure S2:**
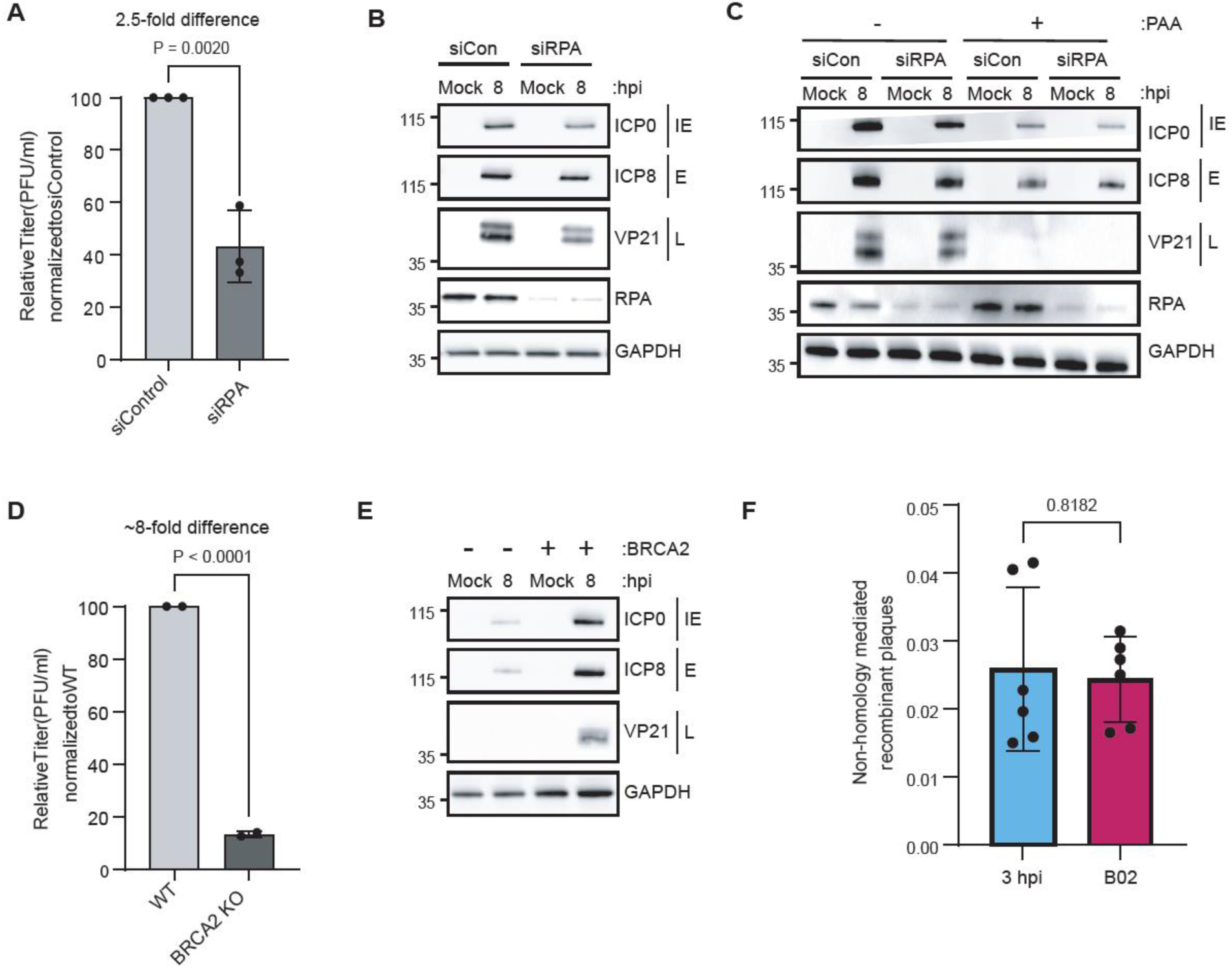
The role of RAD51 during infection is distinct from functions in DNA replication and recombination. **A.** HFF cells were transfected with a control or RPA siRNA. 48 hours post transfection, cells were infected with HSV-1 17syn+ at MOI 0.1 and harvested for plaque assay at 24 hpi. Plaque assay was performed in Vero cells and plaques were counted after 3-4 days. **B and C.** HFF cells were transfected with a control or RPA siRNA. 48 hours post transfection, cells were infected with HSV-1 17syn+ at MOI 3 and harvested for western blot at 8 hpi. C. PAA was added at 1 hpi. **D.** PEO1 WT or BRCA KO cells were infected with HSV-1 17syn+ at MOI 0.1 and harvested for plaque assay at 24 hpi. Plaque assay was performed in Vero cells and plaques were counted after 3-4 days. **E.** PEO1 WT or BRCA KO cells were infected with HSV-1 17syn+ at MOI 3 and harvested for western blot at 8 hpi. **F.** HFF cells were treated with DMSO or 20uM B02 for 2 hours pre-infection. Cells were co-infected with OK11 and OK42 at MOI 20 and Non-homology-mediated recombinant plaques were quantified.

**Figure S3:**
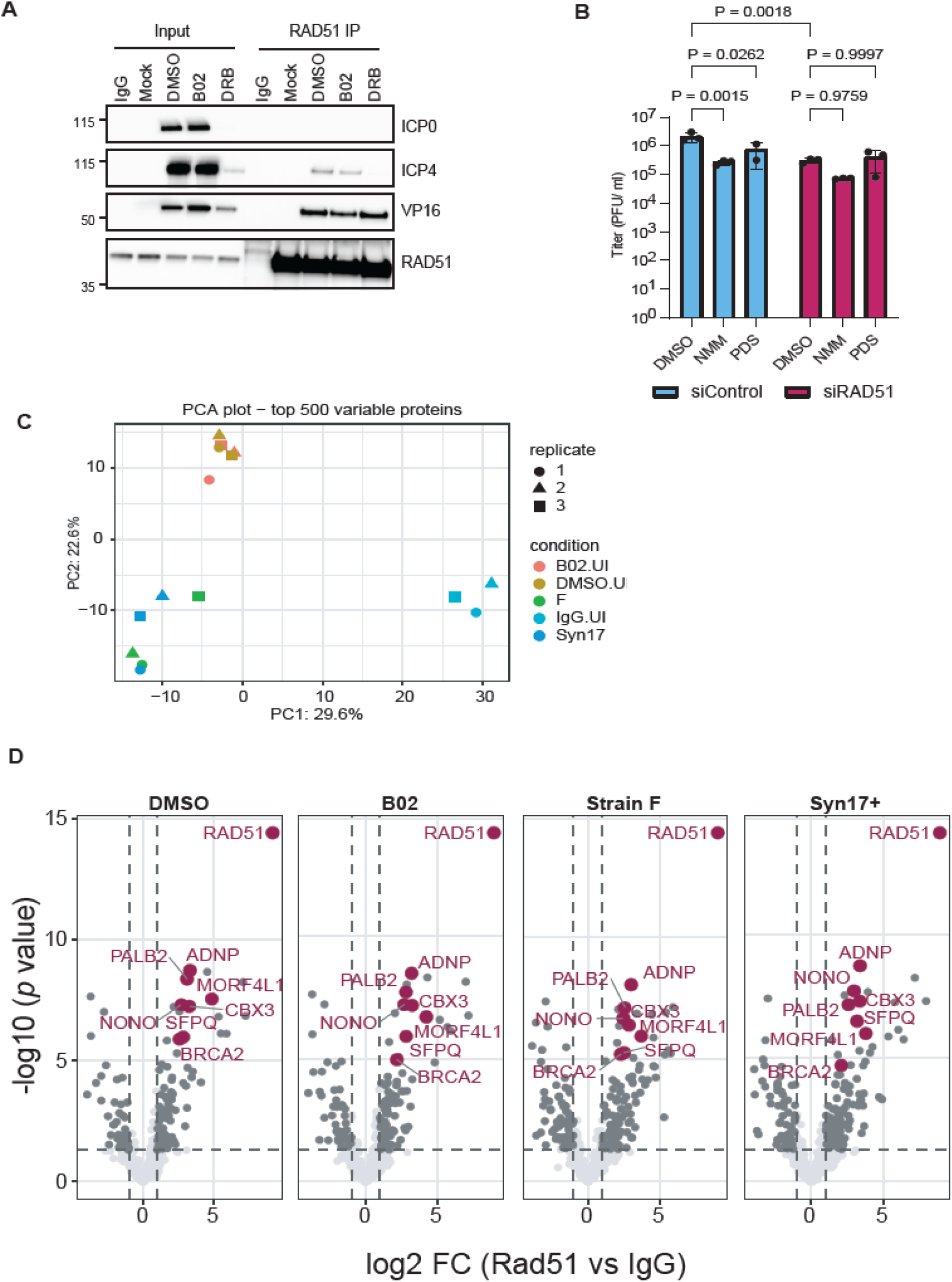
RAD51 interacts with early host and viral transcription factors to and with ADNP: a core factor of the ChAHP complex. **A.** HEp2 cells were infected with HSV-1 17syn+ MOI 10 and RAD51 IP-Wb was performed at 3hpi. Western blot analysis was performed for IE viral proteins ICP0, ICP4 and VP16. Cells were treated with DMSO/ B02 for 16 hours pre-infection and with DRB at 1hpi. **B.** HFF cells were transfected with control, and RAD51. 48 hours post transfection, cells were infected with HSV-1 17syn+ at MOI 0.1 and harvested for plaque assay at 24 hpi. Cells were treated with PDS and NMM at 1hpi. Plaque assay was performed on confluent Vero cells. Statistics: Two-Way ANOVA. **C-D:** HEp2 cells were treated with DMSO/ B02 for 16 hours pre-infection and infected with HSV-1 strain 17syn+ for 3 hours and strain F for 1.5 hours. Cells were harvested for RAD51 Immunoprecipitation followed by mass spectrometry (IP-MS). **C.** PCA plot showing replicates from each treatment condition. **D.** Volcano plots showing fold-change of indicated sample over IgG.

